# Representational drift in barrel cortex is receptive field dependent

**DOI:** 10.1101/2023.10.20.563381

**Authors:** Ahmed Alisha, Voelcker Bettina, Peron Simon

**Affiliations:** Center for Neural Science, New York University, 4 Washington Pl., Rm. 621, New York, NY 10003

**Author notes:** Lead Contact: Simon Peron.

**Keywords:** representational drift, cortical plasticity, barrel cortex

## Abstract

Cortical populations often exhibit changes in activity even when behavior is stable. How behavioral stability is maintained in the face of such ‘representational drift’ remains unclear. One possibility is that some neurons are stable despite broader instability. We examine whisker touch responses in superficial layers of primary vibrissal somatosensory cortex (vS1) over several weeks in mice stably performing an object detection task with two whiskers. While the number of touch neurons remained constant, individual neurons changed with time. Touch-responsive neurons with broad receptive fields were more stable than narrowly tuned neurons. Transitions between functional types were non-random: before becoming broadly tuned neurons, unresponsive neurons first pass through a period of narrower tuning. Broadly tuned neurons with higher pairwise correlations to other touch neurons were more stable than neurons with lower correlations. Thus, a small population of broadly tuned and synchronously active touch neurons exhibit elevated stability and may be particularly important for downstream readout.

## INTRODUCTION

Populations of cortical neurons representing specific sensory^1–4^, cognitive^5–8^, or motor^9^ features exhibit a baseline level of change even in the context of behavioral stability. This ‘representational drift’ typically involves changing responses of individual neurons as well as the addition and removal of neurons to and from the responsive population^10^. Such changing population must nevertheless continue to accurately represent a sensory input or effectively evoke a particular movement^11,12^. One proposed solution the brain could employ to mitigate this problem is through the presence of a subset of neurons with greater stability^13^.

Broadening of receptive fields is a common consequence of sensory processing. Could this be accompanied by greater stability? In mouse primary visual cortex, neurons that are members of ensembles, or groups of highly correlated neurons, exhibit elevated stability^14^. Cortical columns^15^ transform sensory input arriving from the thalamus via a feedforward network from L4 to L2^16^, most prominently resulting in receptive field broadening^17–19^. In primary vibrissal somatosensory cortex (vS1), representation of whisker touch transitions from a distributed population of narrowly tuned neurons in layer (L) 4 to a sparser ensemble-based representation consisting of correlated neurons with broader tuning in L2^20^. A key function of early sensory cortical processing may therefore be to transition from a more unstable population of narrowly tuned neurons in L4 to a more stable population of broadly tuned neurons in L2, providing a more reliable code for downstream readout^13^.

We ask whether the transition to broader tuning and greater pairwise correlations from L4 to L2 is accompanied by greater long-term stability among touch neurons. We re-analyze volumetric two-photon calcium imaging^21^ data from published experiments^20^ in which we monitored touch neurons longitudinally in well-trained mice stably performing an object localization task with two whiskers. We find that neurons with narrower tuning, as well as neurons in L4, show lower levels of stability than more broadly tuned superficial neurons. Neurons with greater pairwise correlations to other touch neurons exhibit especially high levels of stability. We also find that functional type transitions are highly structured: to become broadly tuned neurons, non-responsive neurons usually pass through a stage of narrow tuning. Moreover, neurons rarely change preferred touch whisker or direction. Our work shows that vS1 dynamics are highly structured, with broadly tuned touch neurons in L2 showing substantially lower levels of drift, potentially facilitating downstream readout^13^.

## RESULTS

### Tracking functional types of touch neurons over time

To ask whether representational drift was lower among broadly tuned, superficial neurons, we disaggregated a previously published barrel cortex imaging dataset^20^ in time. Transgenic mice^22^ (Ai162 X Slc17a7-Cre) expressing GCaMP6s across all excitatory neurons were trained on a two-whisker object detection task (**Figure 1A**). On each trial, a pole was presented either within whisking range of the two whiskers or in an out of reach position. After pole withdrawal, a sound cued mice to respond by licking one of two lickports. Responses to the right lickport on accessible pole position trials were rewarded, as were responses to the left lickport on out of reach position trials. Once mice reached stable behavioral performance, three 700-by-700 μm planes spaced 20 μm apart in depth were imaged simultaneously. These three planes were dubbed a ‘subvolume’ and each subvolume was imaged for 50-100 trials after which the next subvolume was imaged. We imaged 5-7 subvolumes per mouse (**Figure 1B**), spanning 300-420 μm total. We started imaging at the layer (L) 1-L2 boundary and continued down to L4. For each layer, subvolumes with at least 750 cells in that layer were used, yielding 2,212 ± 485 L2 neurons per subvolume, (Mean ± SD, n=11 subvolumes across 7 mice), 1,900 ± 426 L3 neurons (n=10 subvolumes across 7 mice), and 2,425 ± 448 L4 neurons (n=12 subvolumes across 7 mice). To ensure enough trials for correct classification of neurons (Methods), we created aggregate sessions for each subvolume from multiple consecutive imaging sessions (**Figure 1C**), spanning 2.9 ± 1.7 days and 166 ± 22 trials per aggregate session. All analyses were performed using these aggregated sessions.

**Figure 1.**
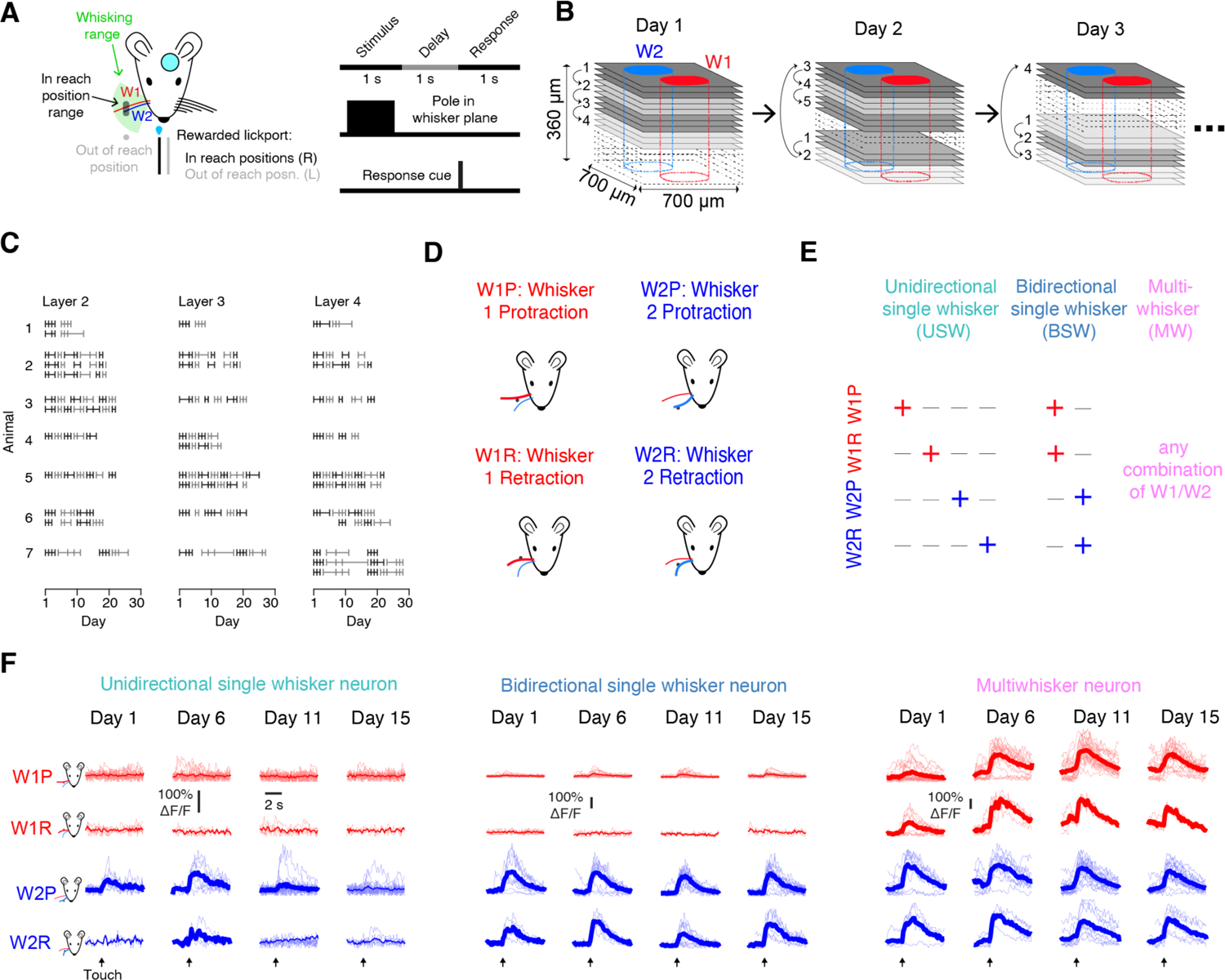
Tracking touch responsive neurons over many days. (A) Behavioral task. Left, trials with a pole appearing in a range of positions (dark grey) accessible to both whiskers are rewarded on right licks. Trials where the pole appears in a position (light grey) outside the reach of whisking (green) are rewarded on left licks. Right, task timing (Methods). (B) Imaging strategy. Groups of three planes (‘subvolume’, same shade of grey) are imaged sequentially on a given day, resuming at the last visited subvolume on the subsequent day. Most but not all subvolumes are typically imaged on any given day. (C) Merging of data across days for each subvolume in the dataset. Vertical lines indicate days for which data was collected; horizontal lines indicate aggregated sessions, grouped to ensure sufficient trials for the encoding model (Methods). (D) Types of touches observed in our task. (E) Classification of neurons into different touch subpopulations. (F) Example neuron responses from a single animal from four time points across two weeks. Light lines, individual trials. Dark lines, mean. Days with significant response to a particular touch type are shown with a thicker mean line.

We used high-speed videography (400 Hz, Methods) to detect and classify touches, using a semi-automated algorithm^23^. Touches were classified (**Figure 1D**) on the basis of the touching whisker (W1, W2, with W2 more anterior to W1) and the direction of touch (protraction, retraction). We used an encoding model that, given whisker curvature change (Δκ), generated a best-possible predicted ΔF/F, incorporating static curvature change kernel and calcium kinetic kernel (Methods). This model measured how well curvature change, a proxy for follicular force ^24^, could predict neural activity. Neurons were considered responsive to a specific touch type if on trials with just that touch type, the encoding model’s predicted ΔF/F response Pearson correlation with actual ΔF/F exceeded 0.15. Neurons that responded to only one whisker-direction combination were considered ‘unidirectional single whisker’ (USW); neurons responding to both directions for a single whisker, ‘bidirectional single whisker’ (BSW); neurons responding to any combination of touches across both whiskers were considered ‘multiwhisker’ (MW; **Figure 1E**). Despite stable touch counts and task performance (**Figure S1**), individual neurons showed varying levels of response stability to individual touch types (**Figure 1F**).

### The size of individual touch subpopulations is stable over time

Representational drift typically takes place in the context of a population of neurons whose size is stable^2^. Before looking at the stability of individual neurons, therefore, we first examined neural stability at the population level (**Figure 2A**). Across all sessions, the average fraction of touch neurons sessions was small, with unidirectional single whisker neurons making up the majority of touch neurons (**Figure 2B**). Because all subvolumes had data to at least day 10, we compared subpopulation fractions from days 1-5 against 6-10. The fraction of neurons that were touch responsive remained stable over the course of imaging (**Figure 2C**; L2, paired t-test, days 1-5 vs. days 6-10, p=0.180, n=11 subvolumes across 7 mice; L3, p=0.347, n=10 subvolumes across 7 mice; L4, p=0.377, n=12 subvolumes across 7 mice). We next measured the fraction of neurons that belonged to each touch subpopulation over the course of all imaging sessions – unidirectional single whisker, bidirectional single whisker or multiwhisker (**Figure 2D**). Because there were so few multiwhisker neurons in layer 4, we excluded these neurons from analysis. The fraction of cells belonging to each subpopulation did not differ between the first five days of imaging and the next five days of imaging in L2 (**Figure 2E**; USW, p=0.158; BSW, p=0.551; MW, p=0.238), L3 (USW, p=0.416; BSW, p=0.537; MW, p=0.135), or L4 (USW, p=0.390; BSW, p=0.055). Thus, across layers, the size of specific touch subpopulations was stable over time.

**Figure 2.**
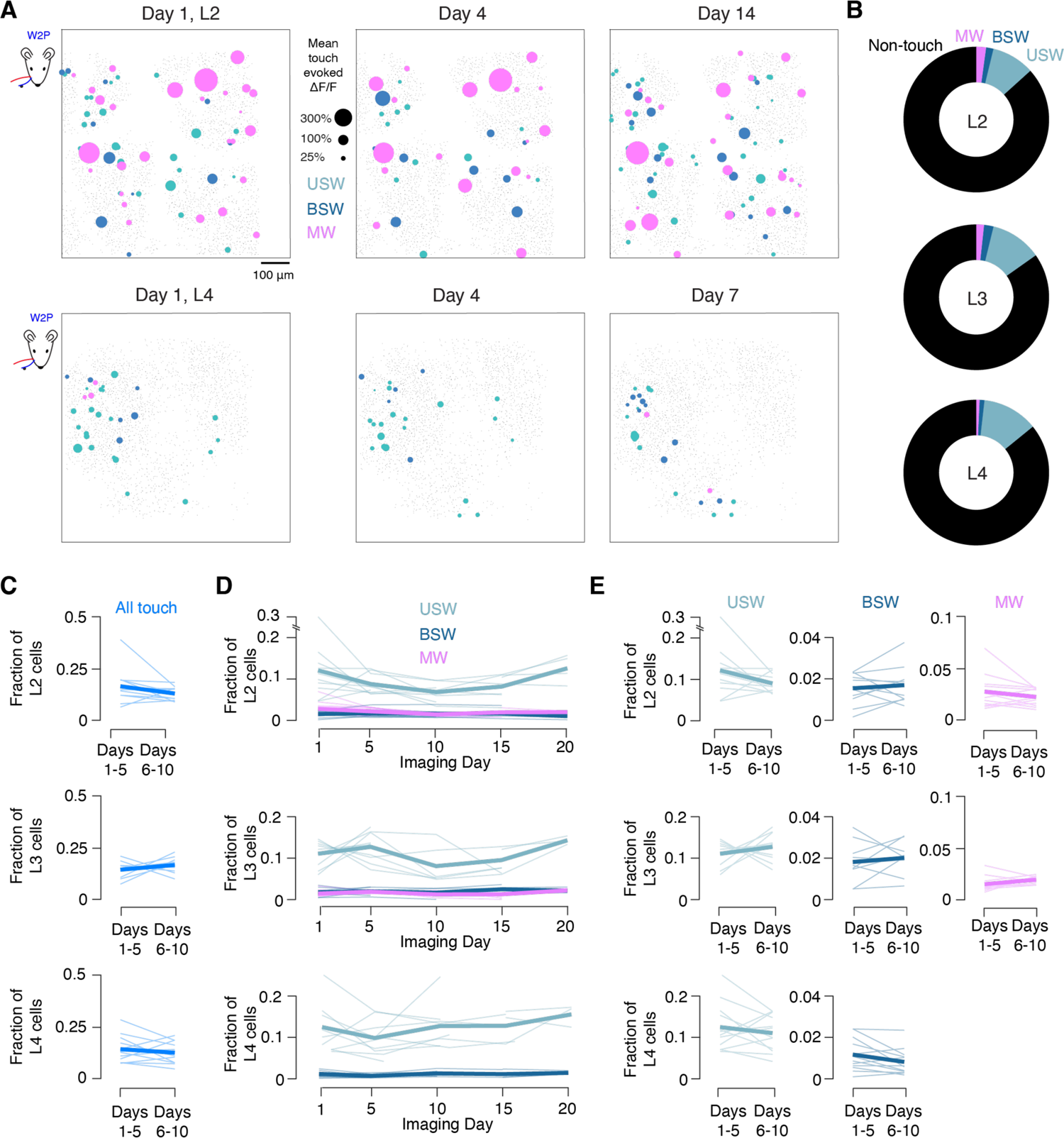
The size of vibrissal representations remains stable over time. (A) Example map showing W2P touch-evoked ΔF/F for unidirectional single whisker, bidirectional single whisker, and multiwhisker neurons across three timepoints for example L2 (top) and L4 (bottom) subvolumes, projected into one plane from three.. Size of circle indicates response magnitude (% ΔF/F) and color indicates subpopulation membership. Light grey points indicate locations of non-responsive neurons. (B) Touch cell subpopulation composition of each layer averaged across all subvolumes and time points. (C) Average fraction of neurons responding to touch on aggregate sessions beginning on days 1-5 and 6-10 for each subvolume in each layer (within-subvolume paired t-test p > 0.05 in all cases). Thin line, individual subvolume. Thick line, cross-subvolume mean. (D) Fraction of neurons in each layer that are unidirectional single whisker, bidirectional single whisker, and multiwhisker across all timepoints for each subvolume in each layer. Aggregate sessions were grouped for days 1-5, 5-10, 10-15 and 20-25, with each multi-day aggregate session assigned to the bin of its first day. Thin line, individual subvolume. Thick line, cross-subvolume mean. (E) Average fraction of neurons belonging to a given touch subpopulation on aggregate sessions beginning on days 1-5 and 6-10 for each subvolume in each layer (within-subvolume paired t-test p > 0.05 in all cases).

### Broadly tuned neurons remain touch responsive for longer periods

Given that we found no change in the aggregate size of the three touch subpopulations, we next compared the stability of the constituent neurons of these populations. We examined neuronal stability by measuring touch responsiveness over time, grouping touch responsive neurons belonging to the same touch subpopulation (USW, BSW, or MW) on the first session of imaging. Restricting our analysis to the more numerous protraction touches, we only considered touch responsiveness to W1P and W2P on subsequent sessions, and only used neurons that were responsive to protraction on the first imaging day. In L2 and L3, neurons that were unidirectional single whisker on the first imaging session showed a larger drop in touch responsiveness at subsequent time points than either bidirectional single whisker or multiwhisker neurons (**Figure 3A**, **B**). This was also observed when the imaging session that started closest to day 5 was used as baseline (**Figure S2A)**.

**Figure 3.**
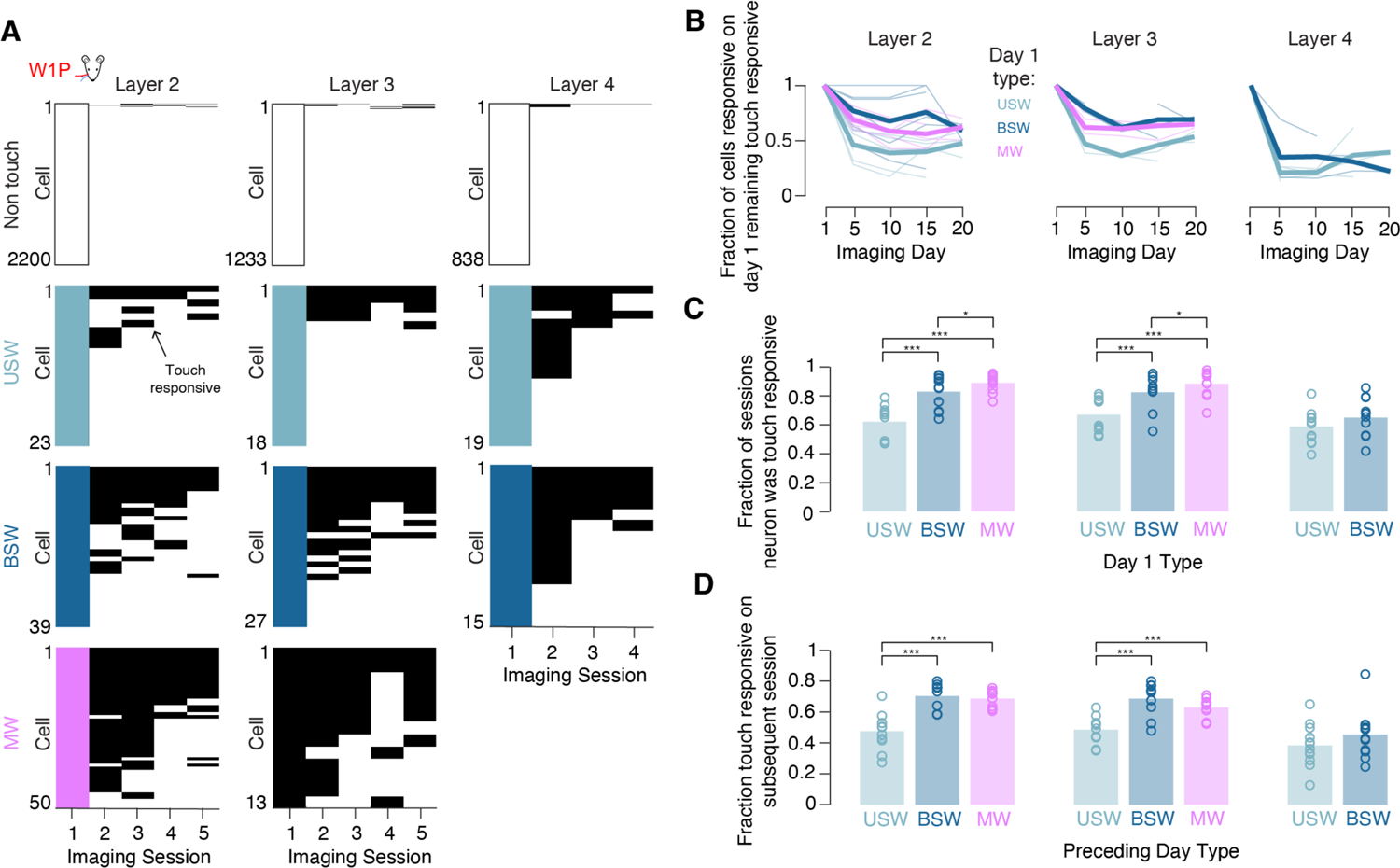
Broadly tuned touch neurons are more likely to remain touch responsive over time. (A) Responsiveness to W1P touches of all neurons over time for an example subvolume from each layer in a single mouse. Black indicates responsiveness on subsequent sessions. Cells are sorted by subpopulation membership on first session and, within subpopulation, on the number of sessions the neuron was touch responsive. Cells that were unresponsive to touch on the first session are shown in white. (B) Fraction of cells that are touch responsive at given time averaged across all mice. Aggregate sessions were grouped for days 1-5, 5-10, 10-15 and 20-25, with each multi-day aggregate session assigned to the bin of its first day. Thin line, individual subvolume. Thick line, cross-subvolume mean. (C) Mean fraction of sessions for which cells of a given subpopulation and layer were touch responsive based on touch subpopulation membership on the first imaging time point. Only cells responsive to W1P or W2P touches on the first session were included, and only responsiveness to W1P or W2P was considered. Bars, mean; circles, individual subvolumes (L2, n=11 subvolumes; L3, n=10; L4, n=12). P-values indicate paired t-test, *p<0.05, **p<0.01, ***p<0.001. (D) Mean fraction of cells remaining touch responsive on subsequent time point given classification on preceding time point. Only cells responsive to either W1P or W2P touches on the first session were included.

To quantify touch responsiveness over time, we measured the average fraction of imaging sessions that neurons belonging to a given touch subpopulation on the first session (USW, BSW, or MW) remained touch responsive. For each neuron in each layer, we calculated the fraction of imaging sessions in which the neuron was responsive to protraction touch (W1P or W2P, calculated separately and averaged; Methods). We then averaged this fraction across neurons belonging to a given subpopulation on the first imaging session. In L2, the fraction of touch responsive sessions was 0.62 ± 0.06 (mean ± standard deviation) for unidirectional single whisker neurons, 0.83 ± 0.06 in bidirectional single whisker neurons, and 0.90 ± 0.03 in multiwhisker neurons (**Figure 3C**), with broadly tuned neurons exhibiting touch responsiveness for more sessions than more narrowly tuned neurons (USW vs. BSW, paired t-test, p<0.001; USW vs. MW, p<0.001; MW vs. BSW, p=0.425). In L3, the fraction of touch responsive sessions was 0.67 ± 0.06 for neurons that were unidirectional single whisker on the first day, 0.82 ± 0.06 in bidirectional single whisker neurons, and 0.88 ± 0.04 in multiwhisker neurons. Again, broadly tuned neurons were more stable (USW vs. BSW, p<0.001; USW vs. MW, p<0.001; MW vs. BSW, p=0.127). In L4, the fraction of touch responsive sessions was 0.59 ± 0.05 for unidirectional single whisker neurons and 0.65 ± 0.06 in bidirectional single whisker neurons, and unlike L2 and L3, the difference was not significant (USW vs. BSW, p=0.134). Thus, the fraction of sessions during which each neuron was touch responsive was higher for broadly tuned neurons compared to unidirectional single whisker neurons in all but L4. When neurons were classified as belonging to a subpopulation on the fifth day of imaging, broadly tuned neurons were still more stable in layer 2 and layer 3 (**Figure S2B**). Unidirectional single whisker neurons were comparably stable across layers. Bidirectional single whisker neurons in L4 were less stable than those in L2 or L3 (L2 vs. L4, unpaired t-test, p=0.002; L3 vs. L4, p=0.004).

We next measured the fraction of neurons of each touch subpopulation that, for any given session, remained touch responsive on the subsequent session (**Figure 3D**). In L2, the average fraction of neurons across animals that remained touch responsive on consecutive sessions was higher in bidirectional single whisker neurons (0.70 ± 0.04; mean ± standard deviation) and multiwhisker neurons (0.69 ± 0.03) compared to unidirectional single whisker neurons (0.60 ± 0.06; paired t-tests: USW vs. BSW, p < 0.001; uSW vs. MW, p < 0.001). Similar trends held in L3 (USW: 0.48 ± 0.04, BSW: 0.67 ± 0.05, MW: 0.63 ± 0.03; USW vs. BSW, p<0.001 USW vs. MW, p<0.001) but not L4 (USW: 0.38 ± 0.06, BSW: 0.44 ± 0.08). Again, unidirectional single whisker neurons were comparably stable across layers, whereas bidirectional single whisker neurons were more stable in L2 and L3 than L4 (unpaired t-test; L2 vs. L4, p=0<0.001; L3 vs. L4, p=0.001). Among multiwhisker neurons, those responding to all four basic touch types were the most likely to be touch responsive on the subsequent session (**Figure S3A**). Among bidirectional single whisker neurons, neurons responsive to the more anterior whisker exhibited greater touch responsiveness on subsequent sessions (**Figure S3B**). Among unidirectional single whisker neurons, retraction preferring neurons exhibited greater responsiveness on subsequent sessions than protraction preferring neurons (**Figure S3C**).

Neurons that were broadly tuned – i.e., multiwhisker or bidirectional single whisker – were thus more consistently touch responsive on subsequent sessions than the more narrowly tuned unidirectional single whisker neurons, with greater stability in L2/3 than in L4.

### Responses of individual broadly tuned touch neurons are more stable over time

Neurons could remain touch responsive over time but nevertheless show a high degree of variability in responsiveness. To quantify neuronal stability more granularly, we computed the correlation of encoding model scores between subsequent imaging session pairs (**Figure 4A**). We computed separate correlations for encoding scores on each protraction touch type (W1P and W2P), averaging for each neuron across the touch type(s) to which it was responsive (i.e., if a neuron responded to W1P and W2P, the average of the two correlations was used, whereas a single correlation value was used for neurons responsive to only W1P or W2P; Methods). Broadly tuned neurons exhibited greater stability than more narrowly tuned neurons, with a decline over time (**Figure 4B**). These trends were also observed when neurons were classified based on their type on the fifth imaging day (**Figure S2C**). We quantified this across layers and touch subpopulations by looking at the mean correlation of encoding scores across consecutive sessions. Correlations were higher in bidirectional single whisker neurons and multiwhisker neurons than in unidirectional single whisker neurons in L2 (**Figure 4C**; USW vs. BSW, paired t-test, p=0.007; USW vs. MW, p < 0.001), L3 (USW vs. BSW, p=0.001; USW vs. MW, p<0.001), and L4 (USW vs. BSW, p=0.046). Encoding score correlations were higher for multiwhisker neurons than for bidirectional single whisker neurons in both L2 (MW vs. BSW, p=0.037) and L3 (MW vs. BSW, p=0.006). Encoding score stability was greater in L2 and L3 than L4 for both unidirectional single whisker (L2 vs. L4, unpaired t-test, p=0.001; L3 vs. L4, p=0.030) and bidirectional single whisker neurons (L2 vs. L4, p=0.003, unpaired t-test; L3 vs. L4, p=0.008). These trends were also observed when neurons were classified based on their type on the fifth imaging day (**Figure S2D**). Among multiwhisker neurons, encoding score stability was lower for neurons responding to more touch types (**Figure S4A**). Encoding score stability was comparable across both whiskers for bidirectional single whisker neurons (**Figure S4B**). Unidirectional whisker neurons showed comparable levels of stability across preferred touch types (**Figure S4C**). In sum, encoding model scores exhibited higher session-to-session correlations in more broadly tuned cells across all layers, and L4 exhibited lower encoding score correlations than either L2 or L3.

**Figure 4.**
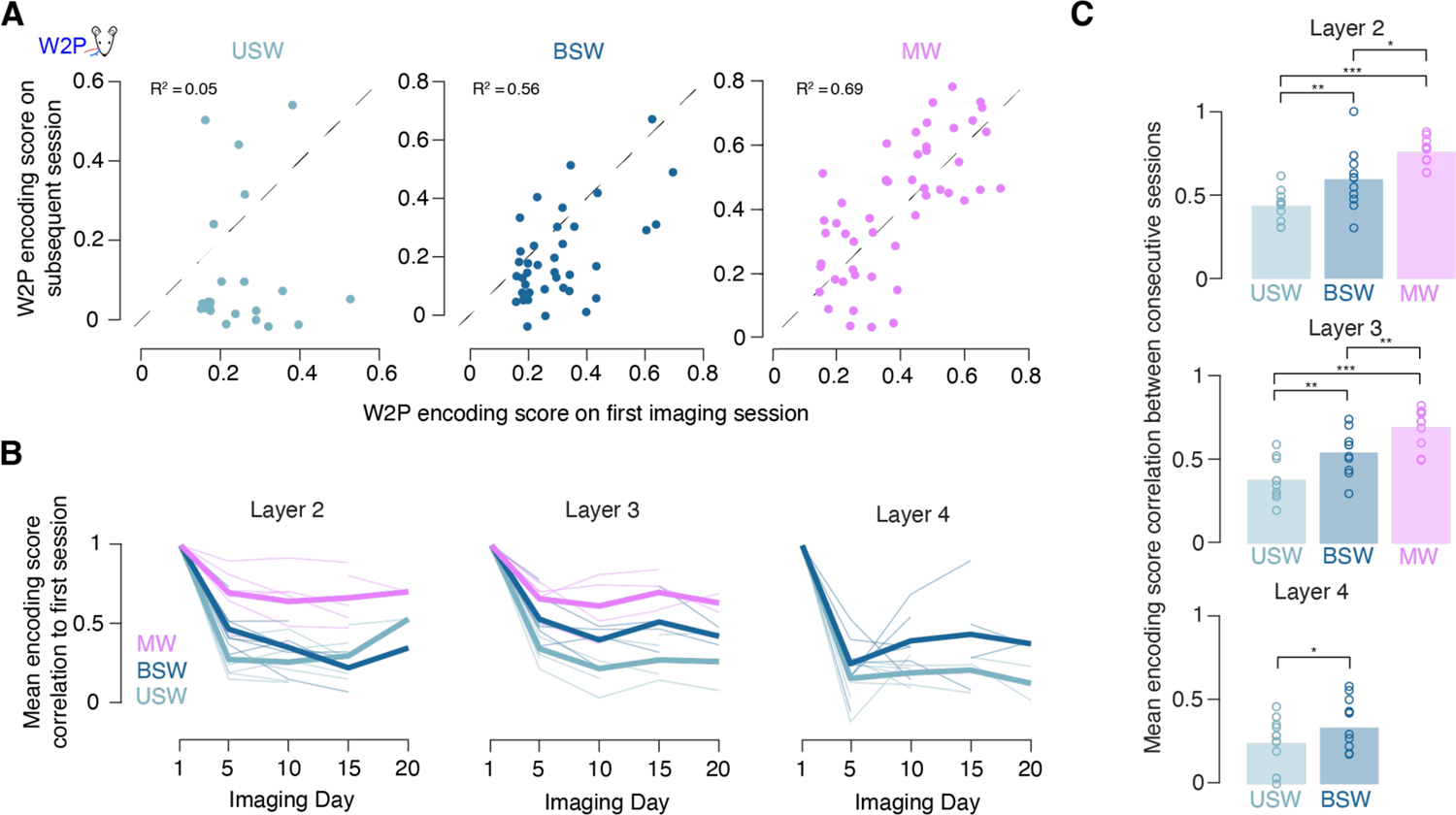
Broadly tuned touch neurons exhibit more consistent responses across time. (A) Encoding scores for W2P touches on the first versus second session for an example mouse for all layer 2 neurons of a given subpopulation that responded to W2P. R^2^ indicates Pearson correlation. (B) Correlation between encoding score on the first session and encoding score on all subsequent sessions. Thin lines, individual subvolumes. Thick lines, mean across subvolumes. Correlations for all touch types for which a neuron was responsive to on day 1 are averaged for subsequent days. Aggregate sessions were grouped for days 1-5, 5-10, 10-15 and 20-25, with each multi-day aggregate session assigned to the bin of its first day. (C) Average Pearson correlation of encoding scores between sessions for all neurons of a given subpopulation and layer, averaged across touch types to which a neuron was responsive. Bars, mean; circles, individual subvolumes (L2, n=11 subvolumes; L3, n=10; L4, n=12). P-values indicated for paired t-tests, *p<0.05, **p<0.01, ***p<0.001.

### Transitions between touch subpopulations are not randomly distributed

We next examined the relative stability of subpopulation membership for individual neurons, looking both at what neurons belonging to a given subpopulation became on subsequent sessions (**Figure 5A**) and which subpopulation a neuron belonged to in prior sessions (**Figure 5B**). We quantified stability by calculating the frequency with which neurons changed touch subpopulation^25^ (**Figure 5C**). Neurons that were non-touch responsive or unidirectional single whisker on a given session were most likely to have been non-touch responsive on the prior session. This was true in all three layers examined (USW probability of having been non-touch on preceding session, L2: 0.58 ± 0.09; L3: 0.59 ± 0.10; L4: 0.69 ± 0.09; non-touch probability of having been non-touch on preceding session, L2: 0.92 ± 0.03; L3: 0.91 ± 0.02; L4: 0.90 ± 0.03).

**Figure 5.**
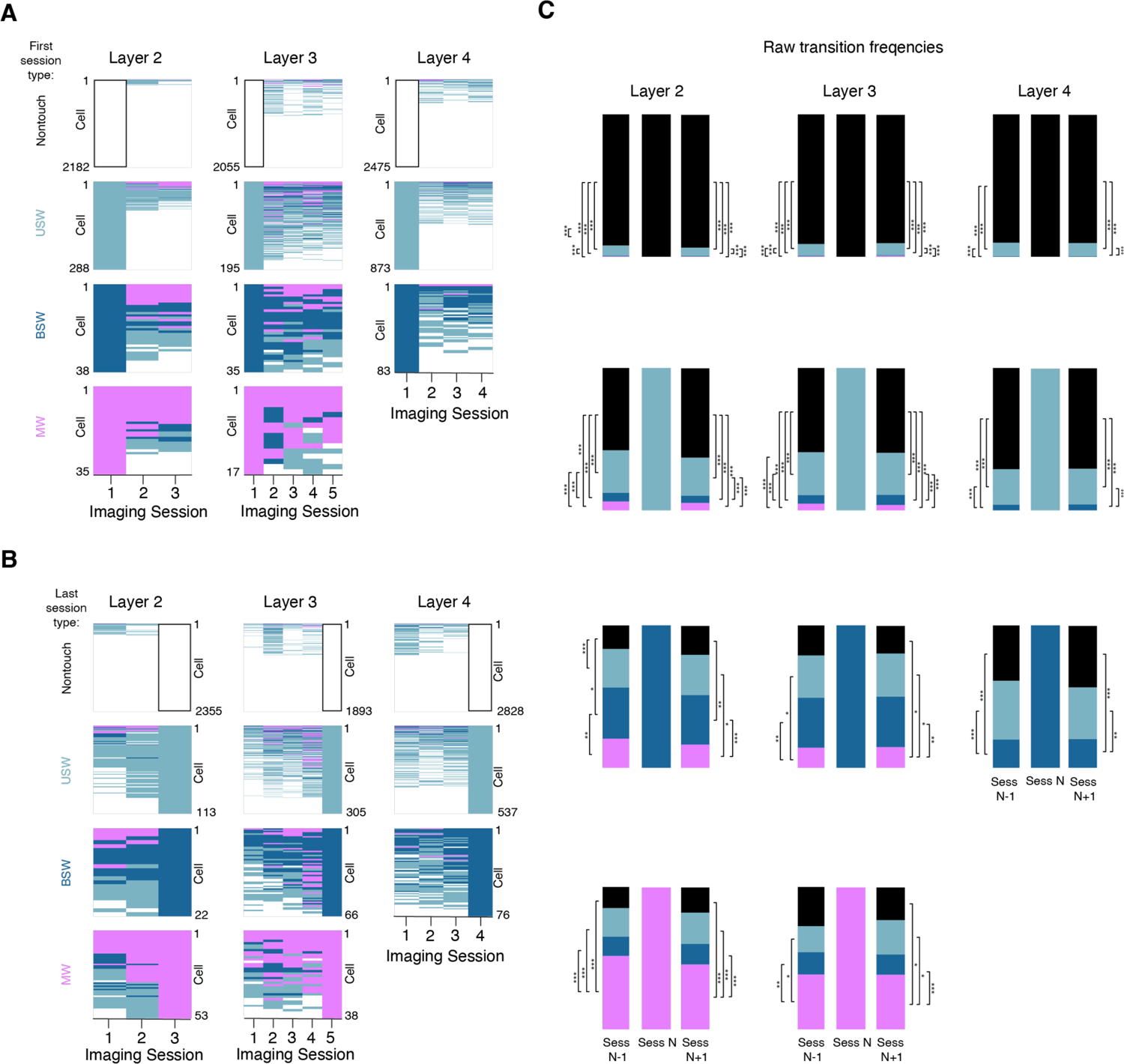
Transitions between neuronal subpopulations are not randomly distributed. (A) Touch subpopulation membership for neurons in example subvolume for each layer, sorted by touch subpopulation (or non-touch, white) on first imaging session. For each cell, its functional type on every subsequent imaged session is indicated. Absence of color indicates non-touch sessions and neurons. Neurons within a subpopulation are sorted by responsiveness. (B) As in A, but grouped based on touch subpopulation on final imaging session. (C) Transition frequencies to and from each neuronal subpopulation for all imaged layers, averaged across subvolumes, sessions and animals. P-values indicated for post-hoc Bonferroni correction following ANOVA comparing composition of all subvolumes within a layer; *p<0.05, **p<0.01, ***p<0.001.

In contrast, for broadly tuned neurons (bidirectional single whisker and multiwhisker neurons), the raw transition frequencies were highest for transitions to and from the same subpopulation. That is, across all layers, bidirectional single whisker neurons had a higher probability of having been bidirectional in the session before and remaining bidirectional the given session than belonging to any other subpopulation. Similarly, multiwhisker neurons in L2 and L3 had a higher probability of having been multiwhisker in the session before and remaining multiwhisker in the session after than of belonging to any other subpopulation. Moreover, bidirectional single whisker neurons had a higher likelihood of having been touch neurons having narrower tuning on the preceding session than of having been non-touch (BSW probability of being non-touch on preceding session, L2: 0.16 ± 0.09; L3: 0.21 ± 0.19; BSW probability of being USW touch: L2: 0.27 ± 0.08; L3: 0.30 ± 0.07; non-touch vs. USW touch, paired t-test, L2: p=0.039; L3: p=0.646). Multiwhisker neurons were also far more likely to have been touch cells with narrower tuning (BSW or USW) than of having been non-touch in both L2 and L3 (MW probability of being non-touch on preceding session, L2: 0.15 ± 0.06; L3: 0.27 ± 0.25; MW probability of being USW or BSW touch: L2: 0.34 ± 0.07; L3: 0.34 ± 0.09; non-touch vs. narrower touch, paired t-test, L2: p<0.001; L3: p<0.001). This implies that the sequence of functional types progresses with higher probability along specific trajectories, with broadly tuned neurons first passing through periods of narrower tuning.

Individual touch subpopulations were different in size (**Figure 2C**), and the raw transition probabilities did not account for this. To assess what transition frequencies look like relative the null model of random transitions between subpopulations, we normalized the raw transition frequencies to the size of each subpopulation (Methods). For all subpopulations except for unidirectional single whisker neurons, normalized transition frequencies were highest for transitions to and from the same subpopulation (**Figure S5**). Normalized transition frequencies for unidirectional single whisker neurons were evenly split between transitions to and from unidirectional single whisker, bidirectional single whisker, and multiwhisker subpopulations. When normalized by subpopulation size, broadly tuned neurons were highly stable – both multiwhisker and bidirectional single whisker neurons were more likely to remain within-subpopulation than to switch.

Whisker preference is stable in L2/3 for neurons preferring the whisker of the barrel they reside in^26^. Was whisker preference also stable in our task? To address this, we tracked whisker preference of neurons across time, finding that neurons were more likely to retain their first-session whisker preference on subsequent sessions than to shift preference (**Figure 6A**, **B**). We quantified this using transition probabilities, restricting analysis only to neurons that were touch responsive on both compared sessions (**Figure 6C**). The probability of retaining whisker preference was high in all layers (probability of whisker 2 tuned cell being tuned to whisker 2 in preceding session: L2: 0.86 ± 0.12; L3: 0.86 ± 0.15; L4: 0.78 ± 0.14; probability of whisker 1 tuned cell being tuned to whisker 1 in preceding session: L2: 0.89 ± 0.12; L3: 0.88 ± 0.09; L4: 0.85 ± 0.09) and significantly greater than the probability of switching (paired t-tests: probability of remaining tuned to whisker 1 vs. probability of switching: L2: p<0.001; L3: p<0.001 L4: p<0.001; probability of remaining tuned to whisker 2 vs. probability of switching: L2: p<0.001; L3: p<0.001; L4: p<0.001). Did directional preference also remain stable? We examined the directional preference of individual neurons over time, also finding it to be stable in individual animals (**Figure 6D**, **E**). Across animals, raw transition frequencies (**Figure 6F**) across directional preferences were low, so that most neurons maintained the same directional preference in addition to whisker preference.

**Figure 6.**
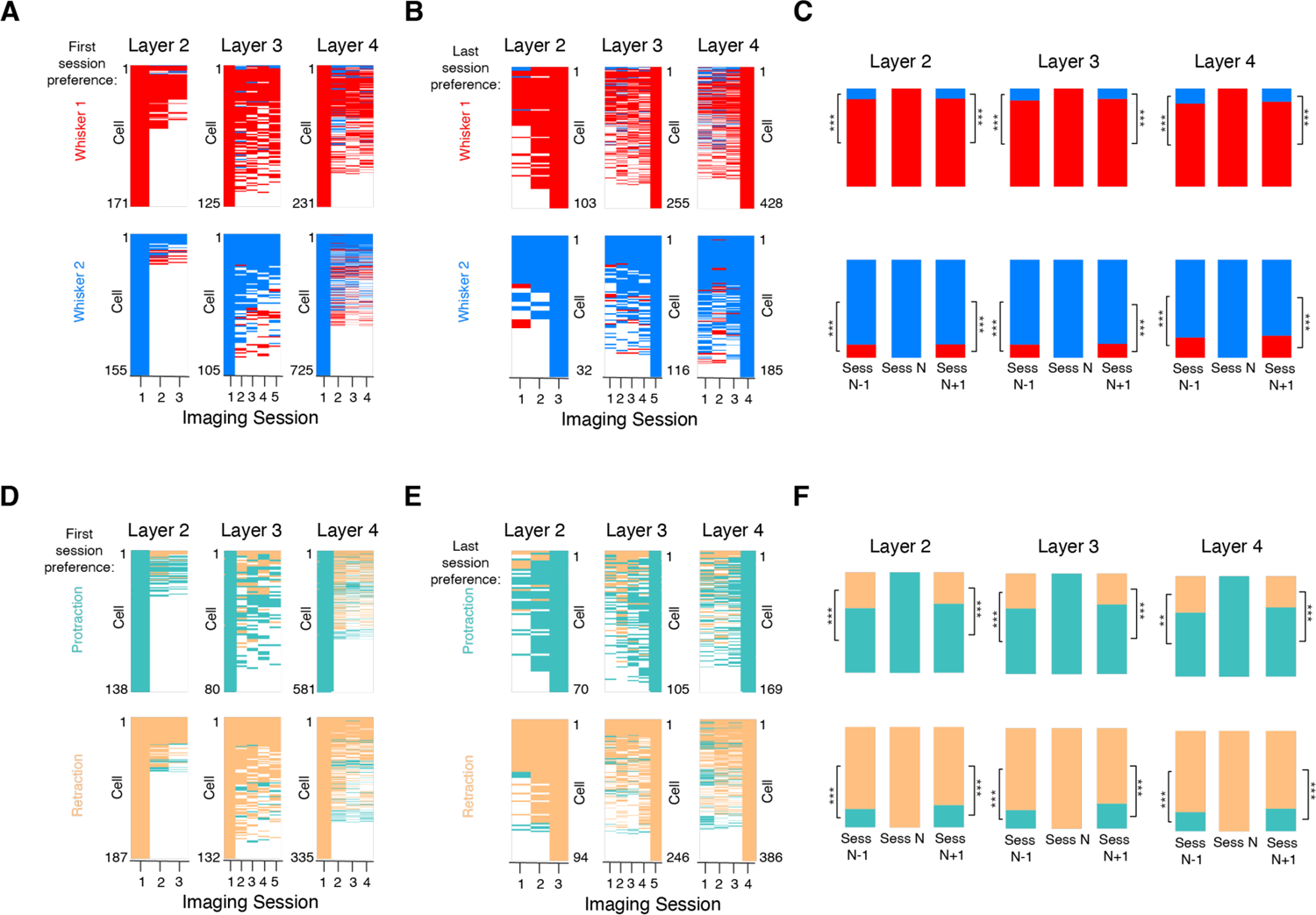
Whisker and direction preference are stable. (A) Whisker preference for neurons in example subvolumes for each layer, sorted by preference on first imaging session. For each cell, its preference on every subsequent imaged session is indicated. Neurons of a given preference are sorted by touch responsiveness. (B) As in A but based on whisker preference on final imaging session. (C) Probability of transition within and between whisker preferences. P-values indicated for within-subvolume paired t-test; *p<0.05, **p<0.01, ***p<0.001. (D) Directional preference for neurons in an example subvolumes from each layer, sorted by preference on first imaging session. For each cell, its preference on every subsequent imaged session is indicated. Neurons within preference are sorted by responsiveness. (E) As in D but based on directional preference on final imaging session. (F) Probability of transition within and across directional preference. P-values indicated for within-subvolume paired t-test; *p<0.05, **p<0.01, ***p<0.001.

Thus, functional type dynamics for touch neurons are non-random. First, broadly tuned neurons are more likely to remain broadly tuned and are more likely to first pass through a stage as more narrowly tuned neurons. Second, neurons rarely switch whisker or directional preference.

### Neurons with higher pairwise correlations are more stable

In L2/3 of mouse visual cortex, members of groups of highly correlated neurons exhibit greater long-term stability to both visually evoked and spontaneous activity^14^. Because elevated correlations in spontaneous calcium signals typically predict connectivity^27,28^, this suggests stability may be a consequence of circuit structure. Do vS1 neurons exhibit similar correlation-dependent stability? To ask this, we examined pairwise correlations during periods without any whisker touch (‘spontaneous’ correlation, Methods) for all touch subpopulations (**Figure 7A**). Among both multiwhisker and bidirectional single whisker neurons, neurons with higher correlations were touch responsive on a greater proportion of sessions (**Figure 7B**). We quantified this by partitioning each touch subpopulation (USW, BSW, and MW) into the top and bottom 25% of neurons based on mean pairwise correlation to other subpopulation neurons. We then computed the fraction of sessions for which neurons in these top and bottom quartiles were responsive. Broadly tuned neurons in the top correlation quartile were touch responsive for longer than narrowly tuned neurons in both L2 (BSW, top: 0.87 ± 0.10, bottom: 0.70 ± 0.14, top vs. bottom quartile, paired t-test, p=0.007, n=9; MW, top: 0.94 ± 0.08, bottom: 0.64 ± 0.12, p < 0.001,n=9) and L3 (BSW, top: 0.84 ± 0.12, bottom: 0.68 ± 0.15, p=0.014, n=8; MW, top: 0.89 ± 0.08, bottom: 0.62 ± 0.13, p=0.001). Unidirectional single whisker neurons only exhibited higher touch responsiveness among more correlated neurons in L3 (USW, top: 0.61 ± 0.19, bottom: 0.45 ± 0.10, p=0.011, n=10), and the difference was smaller than among broadly tuned cells. In L4, correlation did not predict responsiveness (USW, top: 0.49 ± 0.13, bottom: 0.46 ± 0.09, p=0.285, n=12; BSW, top: 0.71 ± 0.14, bottom: 0.66 ± 0.19, p=0.594, n=8). Thus, in L2 and L3, broadly tuned neurons having higher within-subpopulation correlations were more stable than less correlated neurons. This suggests that stability among broadly tuned neurons may be a consequence of elevated connectivity^27,28^.

**Figure 7.**
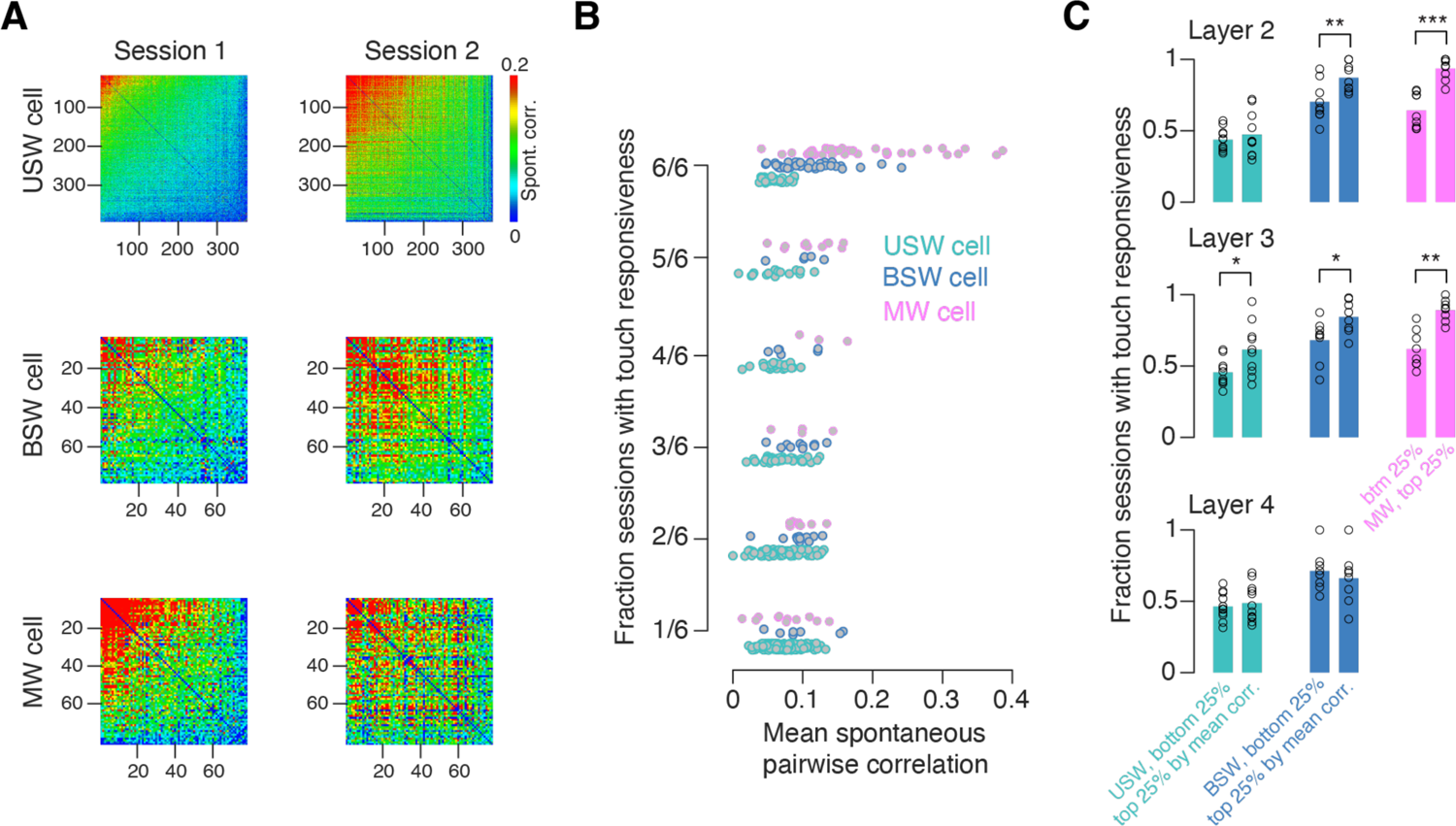
Correlations predict touch responsiveness on subsequent sessions. (A) Pairwise spontaneous activity correlation matrix for an example L2 subvolume for the three touch cell subpopulations on first and second session. (B) Fraction of sessions with touch responsiveness as a function of mean within-subpopulation pairwise correlation for each neuron in an example L2 subvolume. Color indicates touch subpopulation. Fraction of sessions with responsiveness can only take one of six values as there are only six sessions. (C) Fraction of sessions a neuron is touch responsive as a function of pairwise correlations for three different touch subpopulations across all subvolumes and layers. In all cases, the left bar indicates the neurons having the lowest 25% of pairwise within-subpopulation correlations, and the right bar indicates the most correlated 25%. Bar, mean across subvolumes; circle, individual subvolume. P-values indicate paired t-test comparing the top to bottom 25%, *p<0.05, **p<0.01, ***p<0.001.

## DISCUSSION

We find that broadly tuned vS1 touch neurons – bidirectional single whisker and especially multiwhisker cells – exhibit greater stability than more narrowly tuned neurons, with L4 showing lower stability than L2. First, the fraction of sessions during which multiwhisker and bidirectional single whisker cells remain touch responsive is higher than for unidirectional single whisker cells, and in L2/3 than in L4 (**Figure 3**). Second, encoding scores are most stable in multiwhisker neurons and in L2/3, and least stable in unidirectional single whisker neurons and in L4 (**Figure 4**). Third, neurons do not move among different touch subpopulations randomly. Instead, multiwhisker and bidirectional single whisker neurons are more likely to remain within category, unidirectional single whisker neurons are most likely to become non-responsive, and broadly tuned cells typically start out as more narrowly tuned cells (**Figure 5**). Moreover, neurons rarely switch whisker and directional preference (**Figure 6**), implying that certain response features are highly stable. Finally, spontaneous pairwise correlations consistently predict stability of L2/3 but not L4 response within a touch subpopulation over time (**Figure 7**). Together, our results imply that a small population of broadly tuned, highly correlated touch neurons in L2/3 exhibit elevated stability and may comprise a particularly important population for stably representing whisker touch.

Changes in single neuron responses have been observed in the face of stable behavior in primary sensory^1-4,14,26,29,30^, multimodal^7^, and motor cortical areas^31,32^, as well as hippocampus^33,34^. Single neuron responses in vS1 change during whisker-mediated task learning^35–37^ as well as following whisker trimming^38–40^. Even when behavior is stable and sensory input is constant, vS1 touch populations exhibit turnover^2^. We find that broadly tuned neurons – bidirectional single whisker and especially multiwhisker neurons – are more stable than narrowly tuned single whisker neurons. From L4 to L2, representations of whisker touch transition from narrow to broad tuning and from distributed to ensemble based coding^20^. This cross-laminar receptive field broadening is found in many sensory cortices^17–19,41^. Given that neurons participating in ensembles are more stable^14^, a core function of L2/3 may be to transition from less to more stable coding. This is supported by the observation that broadly tuned neurons also dominate vS1 L2/3 output projections to vS2^41^ and vM1^42^. Together, this suggests that less stable neural populations early in the sensory stream funnel activity to more stable populations, which in turn disproportionately impact downstream areas. Such a scheme would both counteract some of the problems posed by unstable representations^11,13^ while allowing earlier stages of processing to benefit from the increased flexibility of unstable responses^10,43^.

Visual cortical neurons did not show layer or areal differences in representational drift, though the laminar trends were consistent with greater L2/3 stability^4^. Though broadly tuned neurons project more prominently to both vS2^41^ and vM1^42^, it remains unclear if those areas exhibit greater representational stability. We did not examine intraday drift due to the low number of trials with specific touch trials. In visual cortex, drift on individual days is largely accounted for by arousal^44^. Pooling across days to generate aggregate sessions should mitigate against such confounds as we pool from a range of times within individual days, and arousal typically varies systematically in individual behavioral sessions.

We show that specific rules govern touch neuron dynamics. To become broadly tuned, neurons typically first pass through a stage with narrower tuning. Given that we do not record continuously, it is likely that the incidence of such transitions through intermediate states is even higher than observed. One explanation for such dynamics is that neurons change their tuning due to long term fluctuations in excitability. Multiwhisker cells in vS1 L2/3 generally show different response amplitudes to various touch types^20,45–47^. If this non-uniformity in input from individual whiskers is static, then as responsiveness increases, responsiveness to specific touch types will appear in a specific sequence, broadening the receptive field. Excitability is governed by many biophysical properties^48,49^; if these fluctuate on the relevant timescale, they could produce the observed dynamics with no changes in underlying synaptic connectivity. Alternatively, fluctuations in excitability could drive plasticity and also produce drift, as observed in hippocampal data^50^. Finally, synaptic plasticity alone could account for the observed changes, with elevated connectivity among broadly tuned neurons acting to stabilize their responses^14^ while narrowly tuned neurons that lack such stabilizing connectivity exhibiting greater instability.

Neurons were unlikely to switch whisker or direction preference. This implies that decoding of touching whisker identity and touch direction is less daunting than it seems: if only 10% of C2 whisker preferring neurons are responsive at any given moment but the entire potentially responsive population is fixed, a decoder need only sample from this C2 superset to ensure consistent readout regardless of which 10% of neurons are active at any given moment. Consistent with this, touch neurons tuned to their barrel’s principal whisker are more stable than neurons tuned to other barrels^26^.

Broadly tuned neurons with the highest pairwise correlations to the other neurons in their subpopulation exhibited the greatest degree of stability (**Figure 7**). In mouse visual cortex, a core subset of ensemble neurons exhibits elevated stability during both spontaneous activity and visual stimulation^14^. Correlated spontaneous L2/3 calcium activity predicts connectivity^28,51,52^. In vS1, declines in touch responsiveness following touch neuron ablation^53^ and channeling of activity towards natural touch responsive populations^54^ point to elevated connectivity among touch neurons. Physically, then, a subset of neurons exhibiting elevated connectivity and greater long-term stability may act as a scaffold for less stable neural activity. The presence of such ‘anchor ensembles’ would facilitate downstream readout while sampling input from more plastic populations that would benefit from greater flexibility, balancing the benefits of a sparse and stable neural code^55–57^ with those of more flexible representations^10,43^.

We find that touch responses are more stable among broadly tuned neurons exhibiting high pairwise correlations with other, similar neurons. Stability – along with the presence of these broadly tuned, highly correlated neurons – increases from L4 to L2. Our results suggest that L2/3 functions to stabilize neural activity by transitioning to an ensemble-based coding scheme with broad selectivity.

## ACKNOWLEDGMENTS

All data were originally collected by Bettina Voelcker^20^. We thank Lauren Ryan for comments on the manuscript. We thank Jayeeta Basu, André Fenton, and Dan Sanes for discussion. This work was supported by the Whitehall Foundation and the National Institutes of Health (R01NS117536).

## AUTHOR CONTRIBUTIONS

A.A., B.V. and S.P. designed the study. B.V. carried out experiments and collected the data. A.A. performed the data analysis. A.A., B.V. and S.P. wrote the manuscript.

## METHODS

### Animals

Adult Ai162 (JAX 031562) X Slc17a7-Cre (JAX 023527) mice of mixed sex^22^, which express GCaMP6s exclusively in excitatory neurons, were used throughout. To suppress transgene expression during development, breeder mice were fed a diet including doxycycline (625 mg/kg doxycycline; Teklad), so that all experimental mice received doxycycline until weaning. All animal procedures were approved by the New York University Animal Welfare Committee.

### Surgery

For cranial window and headbar implantation, each mouse (8-10 weeks old) was anesthetized with isoflurane (3% induction, 1.5% maintenance). A titanium headbar was attached to the skull with cyanoacrylate (Vetbond). A circular craniotomy (3.5 mm diameter) was made in the left brain hemisphere over vS1 (center: 3.3 m lateral, 1.7 mm posterior from bregma) using a dental drill (Midwest Tradition, FG1/4 drill bit). After the craniotomy, a double layer cranial window, which was assembled by gluing a 3.5 mm circular #1.5 coverslip to a 4.5 mm circular #1.5 coverslip (Optical Adhesive 61, Norland Products), was placed over the craniotomy. The cranial window and headbar were further affixed to the skull with dental acrylic (Orthojet, Land Dental).

### Behavior

Following recovery from surgery, mice were trimmed to two whiskers whose barrels had the least obstructive vasculature, typically C2 and C3, and in some cases, C1 and C2. Mice were placed on water restriction, and subsequent trimming occurred every 2-3 days. Water-restricted mice were head-fixed to the behavioral apparatus and trained on an object localization task^20^, in which a metal pole (0.5 mm diameter Drummund Scientific, PA, USA) was vertically moved into the range of the mouse’s whiskers either at an out-of-reach position or at a range of accessible proximal positions. On each proximal trial, the pole appeared at a random position drawn from a range spanning 5 mm along the anterior-posterior axis. In all trials, the pole remained accessible for 1-2 s, after which it was moved downward and out of reach. Pole insertion and removal was accompanied by a 50 ms white noise sound (60-70 dB). 500 ms after the pole was withdrawn, an auditory cue (3.4 kHz, 50 ms) was provided to indicate to the mouse to make a lick response to receive a water reward. Licking the left lickport was rewarded on distal trials, while licking the right lickport was rewarded on proximal trials. On all trials, the lickport remained withdrawn except for during the response epoch (after the auditory cue). Incorrect responses resulted in a timeout (5s) and premature withdrawal of the lickport. Correct trials typically lasted 10 s while incorrect trials lasted 15 s, with mice averaging ∼5 trials per minute. Mice were considered to reach criterion performance once d-prime exceeded 1.5 for two consecutive days, at which point imaging began.

### Whisker videography

Whisker video was acquired using custom MATLAB (version 2019a; MathWorks) software from a CMOS camera (Ace-Python 500, Basler) running at 400 Hz and 640 × 352 pixels and using a telecentric lens (TitanTL, Edmund Optics). Illumination was produced via a pulsed 940 nm LED (SL162, Advanced Illumination). 7-8 s of each trial were imaged, covering the 1 s prior to pole movement, the period while the pole was in reach, and several seconds after the pole was withdrawn. Data was processed on the High Performance Computing (HPC) cluster at New York University. First, candidate whiskers were detected with the Janelia Whisker Tracker. Next, whisker identity was refined and assessed across a single session using custom MATLAB software^23^. Following whisker assignment, curvature (κ) and angle (θ) were calculated at specific locations along each whisker’s length. Change in curvature, Δκ, was calculated relative to a resting angle-dependent baseline curvature value obtained during periods when the pole was out of reach. Next, automatic touch detection was performed. Touch assignment was manually curated using a custom MATLAB user interface. Protraction touches were assigned negative Δκ values.

### Volumetric two-photon imaging

Two-photon calcium imaging was performed using a custom MIMMS two-photon microscope (http://openwiki.janelia.org/wiki/display/shareddesigns/ MIMMS), consisting of a 940 nm laser (Chameleon Ultra 2; Coherent) with power rarely exceeding 50 mW. For imaging of each mouse, multiple subvolumes, each consisting of three 700 x 700 μm imaging planes (512 x 512 pixels) which were spaced 20 μm apart, were acquired at a ∼7 Hz frequency. On a given imaging day, 4-7 subvolumes were imaged sequentially for 50-70 trials each (Figure 1B). After the first imaging day, motion corrected mean images were collected for each plane and used as reference images during experiments on subsequent days to ensure alignment. Depth was modulated with a piezo (P-725KHDS; Physik Instrumente). Laser power was depth-adjusted with an exponential length constant having a value of 250 μm. Imaging data was acquired using Scanimage (version 2017; Vidrio Technologies).

### Imaging data processing

Imaging data was processed on the New York University HPC cluster immediately following acquisition. First, motion correction via image registration was performed. Next, neurons were detected on the first day of imaging using an automated algorithm based on template convolution. The segmentation was verified with post-hoc manual curation. A reference segmentation was thereby established for each plane. On subsequent imaging days, the reference segmentation was algorithmically transferred to the new data^32^. Following segmentation, neuropil subtraction and ΔF/F computation were performed. For layer assignment, each neuron was first assigned a depth in reference to a single reference plane taken at the top of the dura^20^. The L1-L2 border was defined at the depth of the most superficially imaged excitatory neuron and the L3-L4 border was found by manually locating a noticeable shift in neuron morphology in conjunction with the emergence of clearly visible septa. The L2-L3 border was placed at the midpoint between the L1-L2 and L3-L4 borders.

### Imaging session aggregation

Data from multiple imaging days was aggregated to obtain at least 150 trials per aggregate (Figure 1C). This ensured that there were sufficient trials to employ the encoding model, which depends on having at least several unique trials per touch type.

### Encoding model and neural subpopulation classification

We used an encoding model to assess how well whisker curvature change could predict neural activity and, therefore, whether a neuron is a touch cell. The model predicts neural activity (ΔF/F), *rmodel*, for each neuron using:

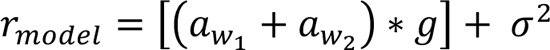

Here, α_wi_ is the predicted amplitude of response to a given whisker’s touch input at a given time, *g* is the GCaMP kinetics kernel, and *σ^2^* is a Gaussian noise term.

For single-whisker touch trials for whisker *i*, α_wi_ is:

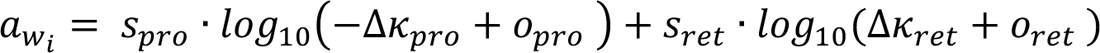

This model is based on previous work using a less constrained generalized linear model that revealed monotonically increasing response as a function of whisker curvature across touch neurons^53^. For a given whisker, the amplitude of the response to a protraction touch (*Δκ* < 0) at a given time, α_wi_, is given by applying a slope *spro* to its change in curvature, *Δκpro*. To account for neurons that have a minimal force needed to elicit a response, the offset term *opro* was included. The retraction (*Δκ* > 0) response is calculated in an analogous manner.

The indicator kinetics kernel, *g*, consisted of a sum of exponentials having time constants *τrise* and *τdecay*. It was normalized so that its peak was 1. Both *τrise* and *τdecay* were constrained based on the known physiological range^58^: *τrise*, 100 ms to 500 ms; *τdecay*,1 s to 5 s. The noise term *σ^2^* was determined for each neuron by measuring the variance of negative ΔF/F values. Our sliding-window F0 fitting procedure^2^, in which we compute F0 using a 3 minute sliding window as the median for neurons that have low activity (non-skewed F distribution) and the 5^th^ percentile for the most active neurons (highly skewed F distribution) ensures that ΔF/F is appropriately 0-centered.

The model was fit with 5-fold cross validation using block coordinate descent and a mean-square-error cost function minimizing the difference between model response, *rmodel*, and neural response, *rneural*. During cross-validation, data was partitioned by randomly drawing 5 disjoint equal sized sets of trials; individual trials were not broken up. The terms of α_wi_ were iteratively fit along with *g* using single-whisker touch trials and an equal number of non-touch trials. Because this resulted in two estimates of *g*, we employed the mean of these parameters for the final model fit.

Neurons were classified based on how well the encoding model could predict neural activity on specific trial types^20^. Neurons were classified as touch responsive for a given session if the correlation between this predicted ΔF/F and actual ΔF/F on a given touch type’s trials was greater than 0.15. Thus, neurons were classified as responsive to each of the four touch types (W1P, W1R, W2P, W2R; Figure 1D) independently. Touch responsive neurons were assigned to touch subpopulations based on their responsiveness to combinations of the four touch types – unidirectional single whisker neurons responded only on one of the four touch trial types, bidirectional single whisker neurons responded to both retractions and protraction of only a single whisker, and multiwhisker neurons responded to any combinations of touch types that involved both whiskers (Figure 1E).

### Touch responsiveness analysis

When performing touch responsiveness analysis over time (Figure 3), only protraction touches were used. For unidirectional single whisker neurons, we only considered a single touch type; for bidirectional and multiwhisker neurons we averaged the responsiveness for the particular touch types (W1P, W2P) that the neurons was responsive to on the baseline day. Thus, for neurons responsive to both W1P and W2P on day 1, responsiveness could take a value of 0, 0.5, and 1. To produce the averaging across imaging days plot (Figure 3B), we binned across days 1-4, 5-9, 10-14, 15-19 and 20-24, based on the imaging day of the first session in an aggregate session.

### Encoding score correlation analysis

For all neurons of a given subpopulation, encoding scores, were calculated for all imaging sessions (Figure 4). For each touch subpopulation (USW, BSW, MW), a vector of encoding scores for both protraction touch types (W1P, W2P) consisting of all neurons that exhibited significant (i.e., greater than 0.15) responsiveness to that touch type was constructed for a given day and the subsequent day (criteria only had to be met on the first of the two compared days). The Pearson correlation was computed for vectors from consecutive days, yielding two correlations per touch subpopulation, one per protraction touch type (W1P, W2P). If a neuron was not significantly responsive on the first day to a particular touch, that value was excluded from both vectors. We then averaged the two correlations together to obtain the overall value for that population for that pair of sessions.

### Transition frequency analysis

To analyze the likelihood that a neuron would change tuning from one imaging session to the next, the probability that a neuron of a given subpopulation would transition to or from another subpopulation was calculated. For all neurons that belonged to a specific subpopulation on the first session (non-touch, USW, BSW, MW), the raw transition frequency in the forward direction was calculated as the fraction of neurons that began in a certain subpopulation and ended up in a different subpopulation in the next imaging session (Figure 5). The raw transition frequency in the backward direction was calculated as the fraction of neurons that belonged to a specific subpopulation (USW, BSW, or MW) on the final session and belonged to a different subpopulation in the previous imaging session. To account for differenced in individual touch subpopulation size, raw transition frequencies were normalized to the size of the subpopulation on a given session. Expected transition frequencies were calculated using the size of the neuron population of a given subpopulation on each session. Normalized transition frequencies were calculated by dividing the raw transition frequencies by the expected transition frequencies (**Figure S5**).

### Pairwise correlation versus responsiveness analysis

To assess whether pairwise correlations predicted responsiveness (Figure 7), we first computed pairwise correlations within specific touch subpopulations on the first day of imaging. Specifically, Pearson correlations of ΔF/F were computed for periods where no touch took place by excluding all time points 1 s prior to and 10 s following a touch. This mostly consisted of trials where the pole was out of reach. For every neuron belonging to any given touch subpopulation (USW, BSW, MW), the mean correlation to all other neurons in that group was computed. Correlations were sorted and neurons within a population were partitioned into the top and bottom 25% by correlation. Touch responsiveness was assessed for all subsequent sessions as described above.

### Statistical analysis

For comparisons across two matched groups, the two-tailed paired t-test was used. Pairing was within-animal. Given the large number of groups in the transition analyses (Figures 5, **S4**), these were compared using a one-way ANOVA with a Bonferroni multiple comparison correction.

## SUPPLEMENTARY MATERIALS

**Figure S1, related to Figure 1.**
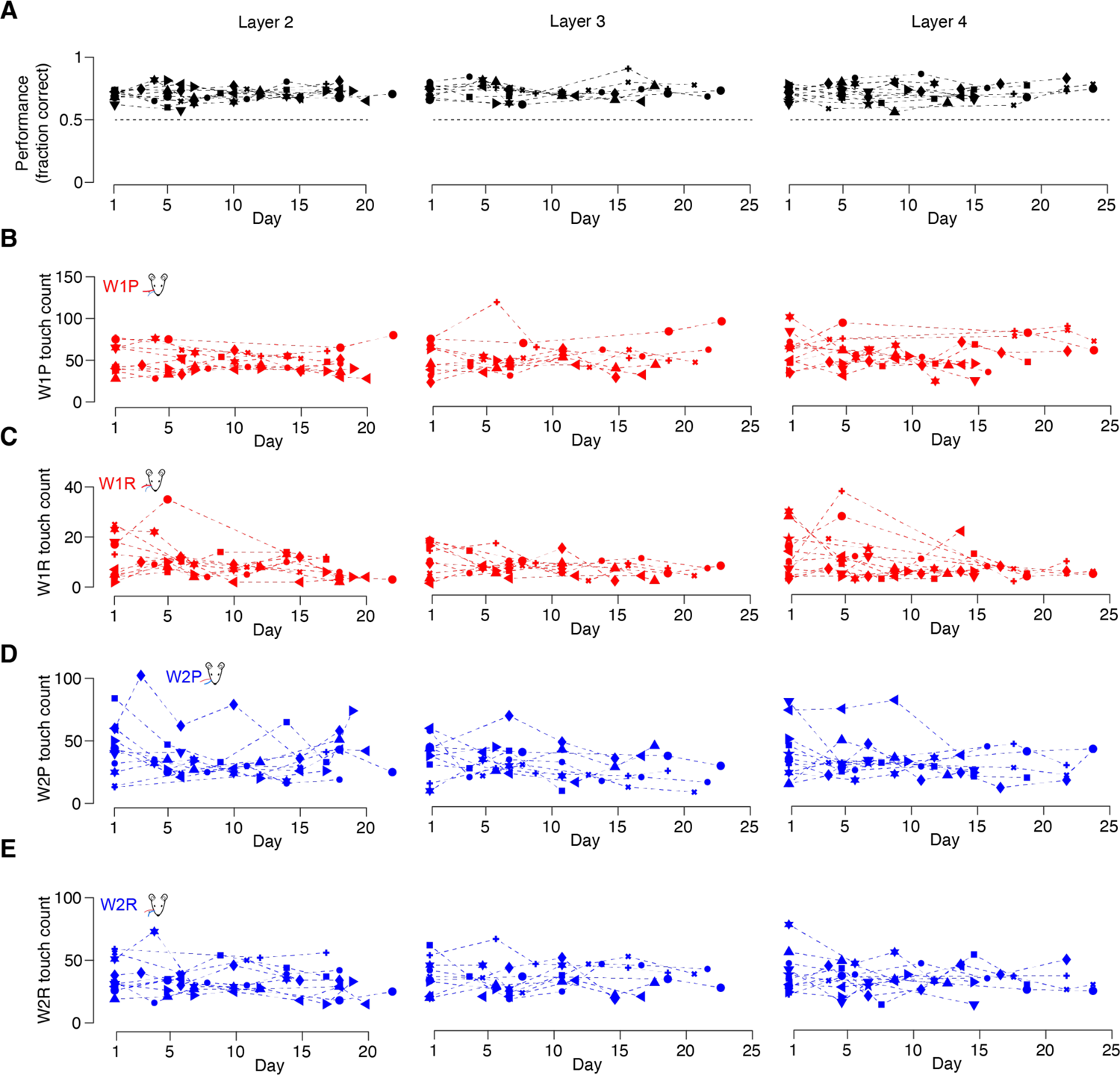
Mice stably perform a two-whisker localization task over weeks. (A) Task performance during each aggregate session block (Figure 1C), with symbol lying on the first day of the aggregate block. Each symbol indicates a specific subvolume. (B-E) Touch counts for the four touch types across all subvolumes and days.

**Figure S2, related to Figure 3 and 4:**
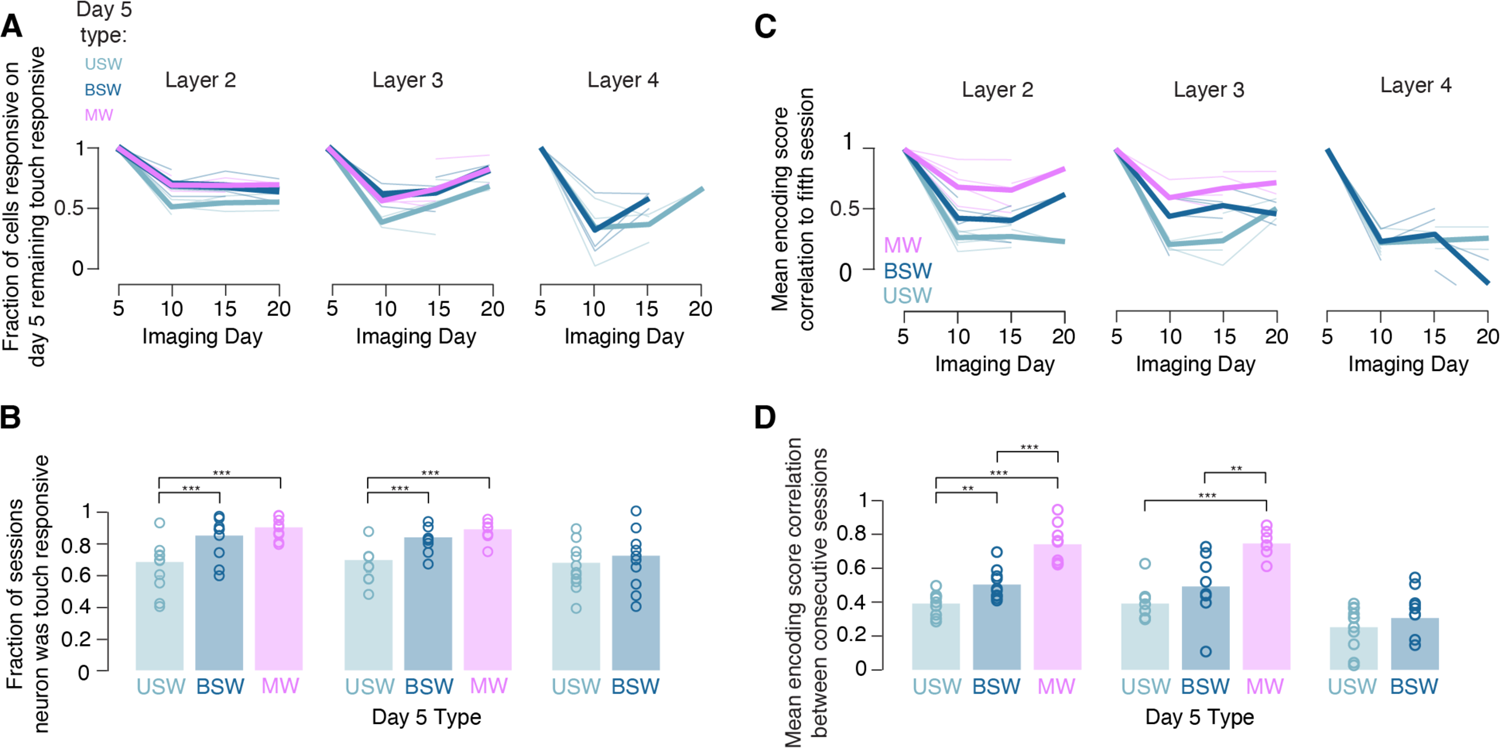
Broadly tuned neurons from day 5 are more stable than narrowly tuned neurons. (A) Fraction of cells that are touch responsive at given time averaged across all mice when classified using day 5. Aggregate sessions were grouped for days 5-10, 10-15, and 20-25, with each multi-day aggregate session assigned to the bin of its first day. (B) Mean fraction of sessions for which cells of a given subpopulation and layer were touch responsive based on touch subpopulation membership on the fifth imaging day. Only cells responsive to W1P or W2P touches on the 5^th^ imaging day were included, and only responsiveness to W1P or W2P was considered. P-values indicate paired t-test, *p<0.05, **p<0.01, ***p<0.001. (C) Correlation between encoding score on imaging session corresponding to fifth imaging day and encoding score on all subsequent sessions. Thin lines, individual subvolumes. Thick lines, mean across subvolumes. Correlations for all touch types for which a neuron was responsive to on day 5 are averaged for subsequent days. Aggregate sessions were grouped for days 5-10, 10-15 and 20-25, with each multi-day aggregate session assigned to the bin of its first day. (D) Average Pearson correlation of encoding scores between consecutive sessions for all neurons of a given subpopulation and layer, averaged across touch types to which a neuron was responsive on the fifth imaging day. Bars, mean (L2, n=11 subvolumes; L3, n=10; L4, n=12). P-values indicated for paired t-tests, *p<0.05, **p<0.01, ***p<0.001.

**Figure S3, related to Figure 3:**
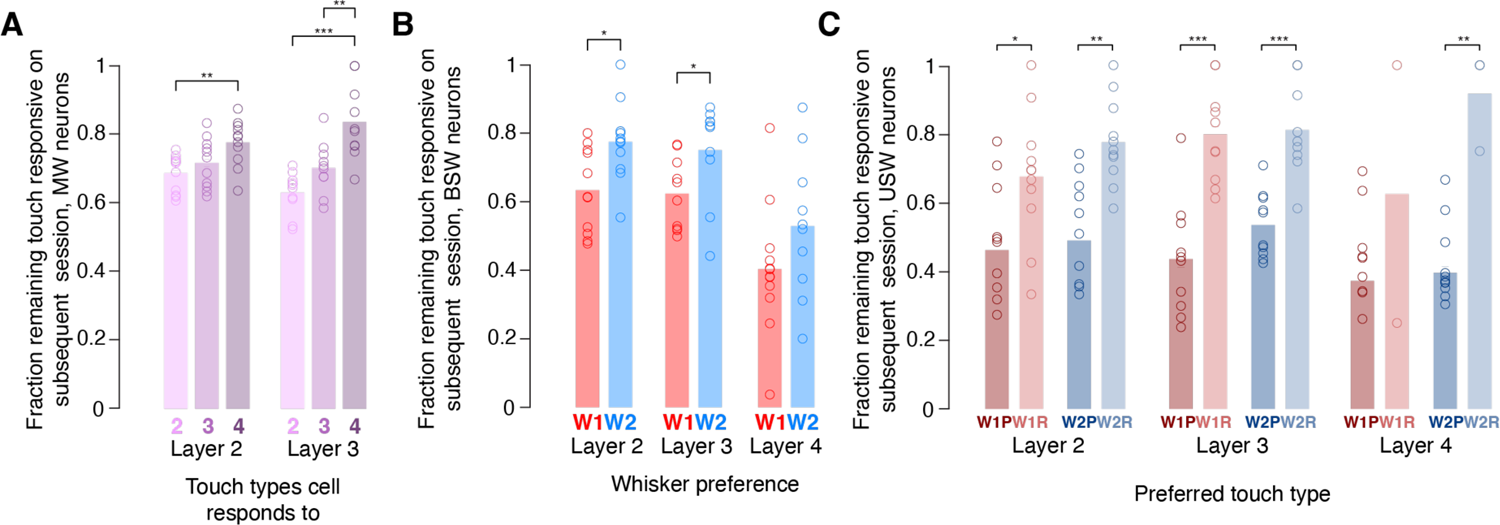
Touch responsiveness over time as a function of receptive field properties of specific touch subpopulations. (A) Mean fraction of multiwhisker neurons remaining touch responsive on subsequent session given the number of touch type primitives (W1P, W1R, W2P, W2R) the cell was responsive to on the preceding session. Bar, mean. L2, n=11 subvolumes; L3, n=10 subvolumes; L4, n=12 subvolumes. P-values indicated for paired t-test comparing subpopulations within-layer, paired by subvolume; *p<0.05, **p<0.01, ***p<0.001. (B) As in A, but for bidirectional single whisker neurons sorted by preferred whisker. (C) As in A, but for unidirectional single whisker neurons sorted by preferred touch type.

**Figure S4, related to Figure 4:**
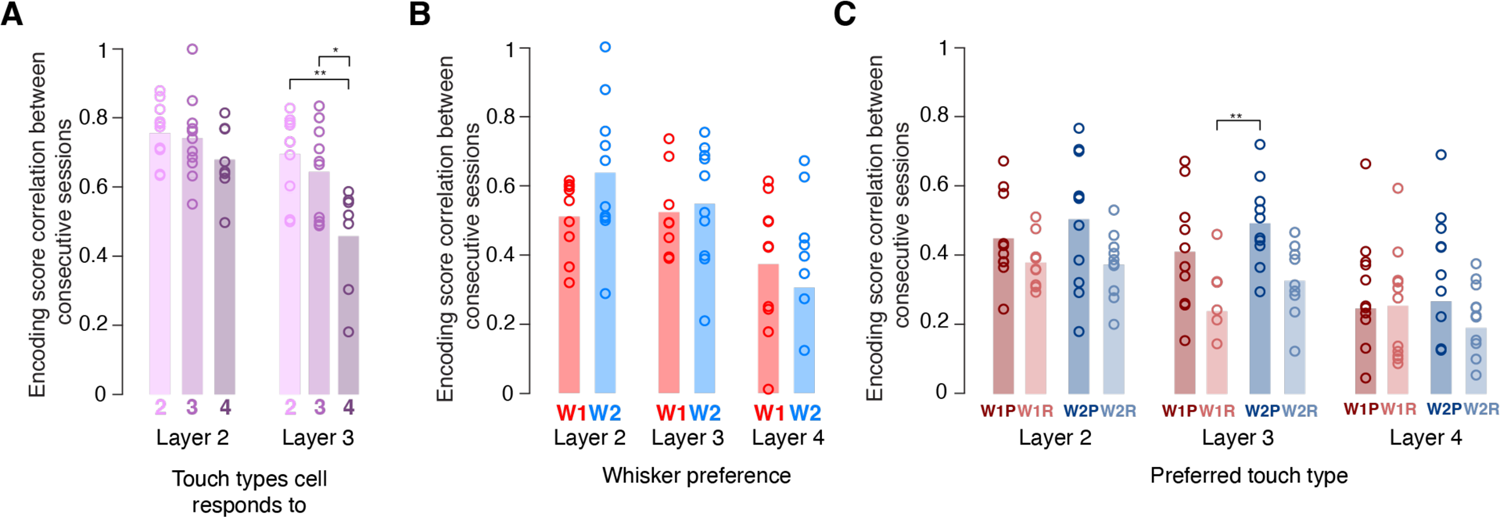
Correlation of encoding score across sequential sessions. (A) Mean Pearson correlation of encoding scores between consecutive sessions for multiwhisker neurons given the number of touch type primitives (W1P, W1R, W2P, W2R) the cell was responsive to on the preceding session. Bar, mean. L2, n=11 subvolumes; L3, n=10 subvolumes; L4, n=12 subvolumes. P-values indicated for paired t-test comparing number of types within-layer, paired by subvolume; *p<0.05, **p<0.01, ***p<0.001. (B) As in A, but for bidirectional single whisker neurons sorted by preferred whisker. (C) As in A, but for unidirectional single whisker neurons sorted by preferred touch type.

**Figure S5, related to Figure 5:**
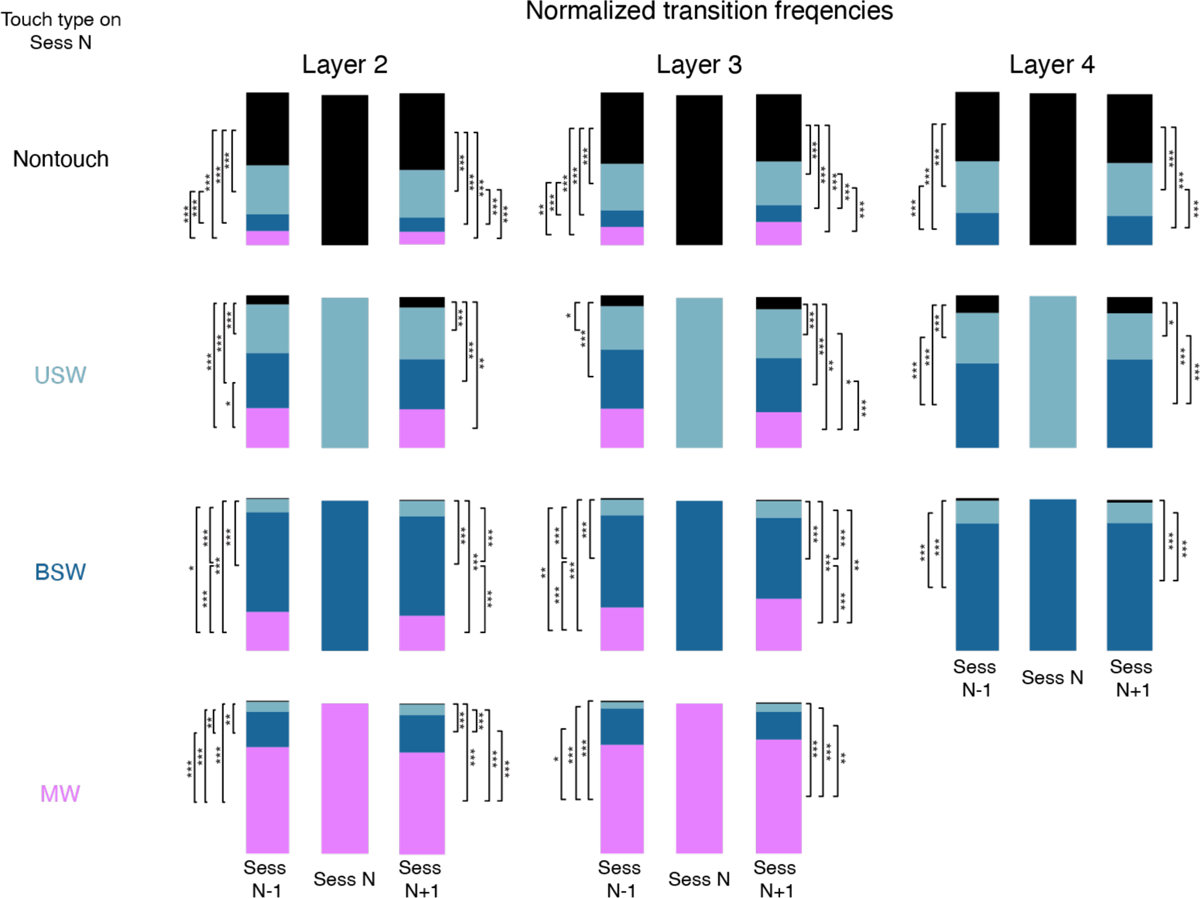
Normalized transition frequencies across touch subpopulations. Transition frequencies of all neuronal subpopulations across layers, averaged across subvolumes, sessions and animals, normalized by size of the population from or to which neurons were transitioning. P-values indicated for post-hoc Bonferroni correction following ANOVA; *p<0.05, **p<0.01, ***p<0.001, comparing all subvolumes within a layer.

